# Embryo development in *Carica papaya* Linn

**DOI:** 10.1101/2021.03.11.434975

**Authors:** Miguel Acevedo-Benavides, Pablo Bolaños-Villegas

## Abstract

Papaya (*Carica papaya* Linn.) is a tropical plant whose draft genome has been sequenced. Papaya produces large fruits rich in vitamins A and C and is an important cash crop in developing countries. Nonetheless, little is known about how the female gametophyte develops, how it is fertilized and how it develops into a mature seed containing an embryo and an endosperm. The Papaya female gametophyte displays a *Polygonum*-type architecture consisting of two synergid cells, an egg cell, a central cell, and three antipodal cells. Reports are available of the presumed existence of varieties in which cross fertilization is bypassed and autonomous development of embryos occurs (e.g., apomixis). In this study, we analyzed the development of female gametophytes in a commercial Hawaiian parental line and in the presumed apomictic Costa Rican line L1. Samples were collected before and after anthesis to compare the overall structure, size and transcriptional patterns of several genes that may be involved in egg and endosperm cell fate and proliferation. These genes were the putative papaya homologs of *ARGONAUTE9* (*AGO9*), *MEDEA* (*MEA*), *RETINOBLASTOMA RELATED-1* (*RBR1*), and *SLOW WALKER-1* (*SWA1*). Our results suggest that its feasible to identify the contour of structural features of *Polygonum*-type development, and that in bagged female flowers of line L1 we might have observed autonomous development of embryo-like structures. Possible downregulation of papaya homologs for *AGO9, MEA, RBR1* and *SWA1* was observed in embryo sacs from line L1 before and after anthesis, which may suggest a tentative link between suspected apomixis and transcriptional downregulation of genes for RNA-directed DNA methylation, histone remodelers, and rRNA processing. Most notably, the large size of the papaya embryo sac suggests that it could be a cytological alternative to *Arabidopsis thaliana* for study. Significant variation in embryo sac size was observed between the varieties under study, suggesting wide differences in the genetic regulation of anatomical features.

## Introduction

*Carica papaya* Linn is a diploid species with male, female, or hermaphrodite flowers on the same plant. In nature, plants show male and female flowers on separate individuals (Geetika *et al*., 2018). However, commercial papaya cultivars show hermaphrodite flowers in some plants and only female ones in others (Geetika *et al*., 2018). Papaya is one of the top 10 most important tropical fruit crops in the world, and its consumption has been recommended to prevent vitamin A deficiency in tropical and subtropical developing countries (Liao *et al*., 2017). Nonetheless, truly little is known about reproductive mechanisms involved in papaya sexual and/or asexual reproduction.

In flowering plants such as maize and *Arabidopsis thaliana*, a megaspore mother cell (MMC) undergoes meiosis to produce four recombinant haploid cells, the megaspores (Rodríguez-Leal and Vielle-Calzada, 2012). Then a single megaspore may give rise to the female gametophyte. After cellular enlargement, the nucleus of the megaspore will undergo three rounds of mitosis to form a gametophyte containing seven cells: two synergids, the egg cell, a binucleated central cell and three antipodals (Rodríguez-Leal and Vielle-Calzada, 2012). Double fertilization of the egg and central cell with one sperm nucleus each initiates the development of embryo and endosperm, respectively (Schmidt *et al*., 2014). In contrast, in clonal species of genus *Boechera*, one can observe 1) regular gametophytic apomixis, in which a diploid sporophytic cell of the nucellus develops in proximity to the MMC and then develops as an embryo or 2) diplospory, a type of gametophytic apomixis, in which the MMC itself becomes an apomictic initial cell that skips meiosis (apomeiosis) and gives rise to an unreduced embryo sac (Schmidt *et al*., 2014; Garman *et al*., 2019). The egg cell then develops into an embryo without fertilization (parthenogenesis). Endosperm development can be autonomous or require fertilization (pseudogamy) (Schmidt *et al*., 2014).

In her seminal article from 1943, Lois Thomson Foster described for the first time the reproductive organs of papaya (Thomson Foster, 1943). In most flowering plants, the development of embryo sacs displays a *Polygonum*-type architecture, an architecture that consists of seven cells: two accessory cells called synergids, which are important for pollen tube attraction; an egg cell, which gives rise to the embryo; a diploid central cell which gives rise to the endosperm; and three cells of unclear function called antipodal cells (Sundaresan and Alandete-Saez, 2010). However, this original work did not detail any of those features; instead, it included very approximate diagrams of tissue sections featuring the integuments, the purported location of the micropyle, the funicle, vascular bundles and the tiny megaspore.

Considering that the draft papaya genome has been published (Ming *et al*., 2008), there should be focus on the development of cellular and molecular tools for studying the reproductive biology of such an important crop. Most notably, reports exist of asexual development of fruits in papaya (Vegas *et al*., 2003). Apomixis has been described as a process that involves development of embryos from sporophytic cells due to changes in the activity of cell-cycle regulators and perhaps due to the activity of *Polycomb*-group proteins that control high-order chromatin architecture (Rodrigues *et al*., 2010). Detailed cytoembryological of embryo sac development should be previously carried out to be able to produce solid information on the reproduction molecular control in this species.

This study aimed to tackle those problems and investigated 1) whether papaya embryo sacs are *Polygonum*-type, 2) whether any structural or developmental variation exists between different lines, and 3) whether any differences exist in the transcriptional patterns of putative regulators of embryo-sac development genes. Our results indicate that it might be methodologically feasible to study embryo sac development, that lines exhibit differences in the size of embryo sacs, that asexual development of embryos may occur in line L1, and that differences in transcriptional patterns occur.

## Materials and Methods

### Plants

Young plants from papaya parental lines L1 (pistillate) and Hawaiian (hermaphrodite) were grown at the Fabio Baudrit Agricultural Research Station at La Garita (San José district), Alajuela province, Costa Rica. The location is 010°00′18″N 084°15′56″W and 840 m above sea level. Average temperature is 22 °C. Plants from were grown under greenhouse conditions with drip irrigation. Each homozygous line was grown separately on dedicated greenhouses and cross pollinations were not performed during sample collection. Flowers were bagged with nylon fabric and collected considering that the time from flower bud emergence to anthesis is 45 to 47 days in pistillate flowers, whereas hermaphrodite flowers show a 2-day delay (Fisher, 1980). Also, because maximum anthesis occurs from 18:00 to 20:00 h (Fisher, 1980), flowers were collected in the early morning of the next day.

### Preparation and analysis of embryo sac sections

Fresh tissue samples (at least three flowers) were fixed in FAA (10% formaldehyde 37% v/v, 5% glacial acetic acid, 85% ethanol 70% v/v) for 24 h at room temperature. Then, samples were dehydrated in a graded series of ethanol (70, 80, 90, 95 and 100%) v/v, for 10 min each, then heated in a microwave to 50-60 °C (45 sec, 100 mL sample). After heating, samples were clarified in two changes of xylene for 10 min, then heated again to 50-60 °C, and finally embedded in Paraplast wax. The resulting samples were cut to 5 µm by using a Reichert-Jung 820H Histostat rotary microtome (Leica, Wetzlar, Germany) and Drukker VC186200 diamond knives (Cuijk, The Netherlands). The samples were dewaxed, hydrated, and stained with nuclear stain Safranin-O (0.5% m/v in 50% v/v ethanol) for 30 min then with membrane stain Alcian blue (0.5% m/v in 3% v/v acetic acid solution) for 30 min. Finally, samples were dehydrated in a graded series of ethanol (95, 100 and 100% v/v), clarified with two changes of xylene, and finally mounted with Permount® (Kurczyńska *et al*., 2007; El Maâtaoui *et al*., 1990). Slides were examined under an Olympus BX53 microscope coupled to a DP73 camera (both from Olympus Corp., Shinjuku, Tokyo). Image cropping and color correction involved using Adobe Photoshop 2020 (Adobe Inc., San Jose, CA, USA).

### Transcriptional analysis

Extraction of mRNA involved the ReliaPrep RNA Cell Miniprep Kit (Promega, Madison, WI, USA), as per the manufacturer’s instructions. RNA quality was evaluated with Qubit 4 fluorometer (Invitrogen, Waltham, MA, USA), and overall integrity was evaluated on 2% agarose gel. Reverse transcription (RT) involved the Access RT-PCR System (Promega), as per the manufacturer’s instructions. Each RT reaction was performed with 31 µL distilled water, 1 µL DNTP, 1 µL each primer, 2 µL MgSO_4_, 46 µL Master Mix, 1 µL polymerase and 2 µL mRNA. Each QPCR reaction was performed with 2 µL SYBR Green, 100 µL GoTaq® qPCR Master Mix, 18 µL nuclease-free water, 40 µL cDNA (20 ng/µL), and 40 µL each primer (150-200 nM) for a total volume of 150 µL. Reactions were run and analyzed on a Rotor-Gene Q system operated with QRex software for comparative quantitation (both from Qiagen, Hilden, Germany). Melting curve analyses and/or negative controls were used to rule out primer-dimer artifacts and low specificity in the amplification reaction. Quantitative reactions were performed in triplicate and averaged. Primers biased for the 3′-end of coding sequences were designed with Primer Express 3.0 and manufactured by Macrogen (Seoul). At least three flowers per line were used.

## Results and Discussion

During the isolation of samples, we found differences in the relative number of ovules per flower (after anthesis). In Hawaiian papaya, we could count approximately 622 ± 15 ovules per flower (*n*=20), whereas in L1, we could count only 140 ± 4 (*n*=20). Sample mass was different as well: after anthesis, the mean weight of ovules per flower in Hawaiian papaya was 874.5 ± 0.3 mg (*n*=20), whereas in L1 flowers, the mean weight was 420.1 ± 0.2 mg (*n*=20). These results suggest that in Hawaiian papaya, each embryo sac has an approximate weight of 1.4 mg, whereas in L1, a single embryo sac may weigh 3.0 mg, so an L1 embryo sac may weigh twice on average.

In Hawaiian ovules (collected 24 h after anthesis), the width was approximately 150 ± 13 µm and length 300 ± 17 µm (*n*=10), whereas in line L1, the width was approximately 310 ± 22 µm and length 530 ± 40 µm (*n* =15), which suggests large natural variation. In contrast, ovules in *Arabidopsis* have an approximate width of 100 µm (Johnson *et al*., 2019). Two possible sources for this variation within papaya may be cell size and cell number. Tissue cells of L1 appear large, with nuclei deeply stained with Safranin-O. Cell numbers per layer appear to vary as well, especially in the inner integument; for instance, Hawaiian papaya has 6±1 cells versus 12±1 cells in L1.

We could distinguish apparent early embryos in both lines (Figure 1C and 1F). No line showed supernumerary or ectopic structures. Flowers collected from line L1 were all pistillate and were bagged 48 h before collection, moreover plants from L1 are routinely grown in a separate greenhouse to prevent pollen contamination. Flowers of the Hawaiian line were not emasculated since removal of anthers causes dehiscence and because it is assumed that only sexual seed is produced. Therefore, only line L1 showed apparent asexual development of embryos. Unfortunately, sample collection did not continue after anthesis; therefore, we could not determine know whether these suspected asexual structures might continue to develop into fully functional seeds that germinate. Asexual development of embryos in L1 may have been overlooked in the past because the plants are continuously used as obligate pollen recipients for breeding of the Costa Rican hybrid cultivar “Pococí”.

**Figure 1.**
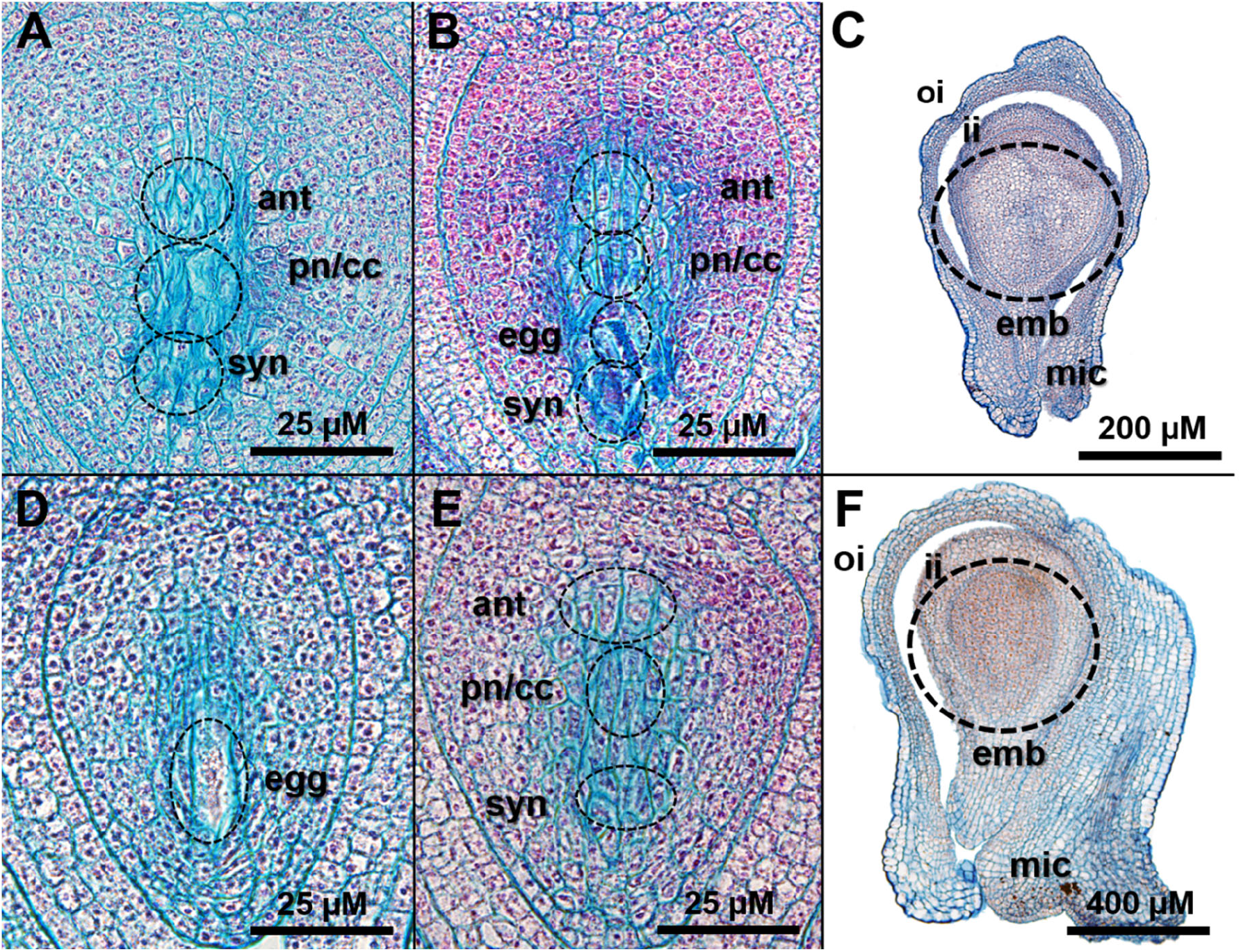
Embryo sac development in papaya lines Hawaiian and L1. Tissue sections of papaya ovules from hermaphrodite, pollen donor line Hawaiian (A-C) and pollen receptor, staminate line L1 (suspected apomictic, D-F), before and after anthesis (24 h). Abbreviations: antipodal cells (ant), central cell (cc), egg cell (egg), embryo (emb), inner integument (ii), outer integument (oi), micropyle (mic), synergid cells (syn), polar nuclei (pn). Line Hawaiian: A) top to bottom, approximate location of the antipodals, central cell (with a seemingly large vacuole stained by Alcian blue) and synergids; B) top to bottom, approximate location of antipodals, central cell, egg cell and synergids, with their nuclei appearing stained red by Safranin; C) developing embryo: the inner and outer integument, micropyle and contour of a nascent embryo can be identified. Line L1: D) ovule showing the approximate location of the egg cell; E) approximate location of the antipodals, polar nuclei and synergids; F) ovule showing a developing embryo, probably of asexual origin, the approximate location of the inner and outer integuments and the micropyle.

Analysis of transcription by qPCR involved the use of primers biased for the 3’ end of coding sequences from the putative homologs of known *Arabidopsis* regulators of embryo sac development (Shi and Yang, 2011) (Table 1). Because no functional validation has been performed yet, the target loci were chosen entirely on the similarity of the papaya sequences to already characterized genes from *A. thaliana* (Table 1). All papaya homolog sequences were significantly similar and included the same functional motifs.

**Table 1.**
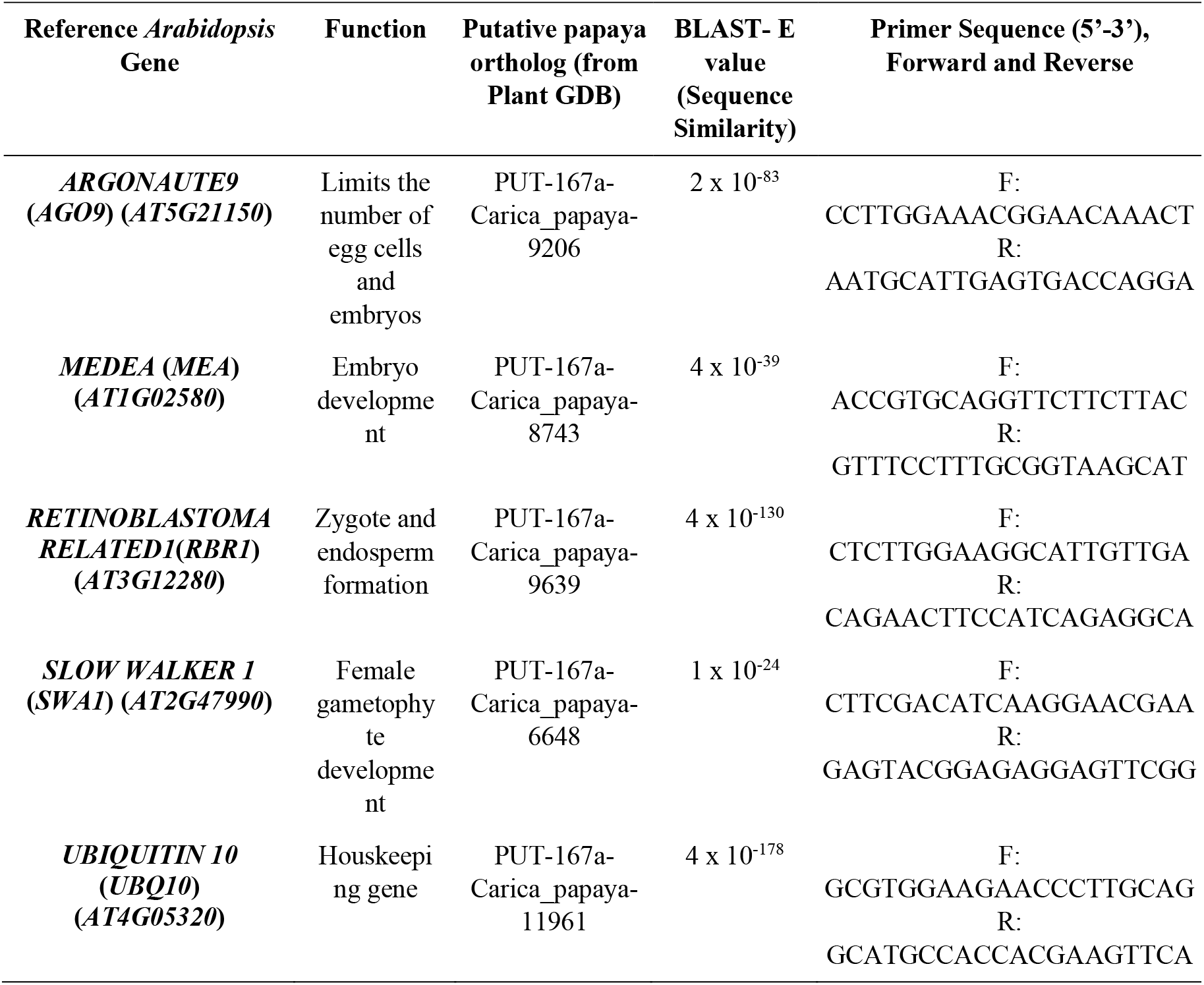
Putative papaya ovule developmental regulators analyzed by quantitative real-time PCR (qPCR). *Arabidopsis* reference genes were selected from relevant sources (Yang et al., 2010; Shi and Yang, 2011). Information was collected from The Arabidopsis Information Resource (TAIR, https://www.arabidopsis.org/), the Plant Comparative Genomics site (Plant GDB, http://www.plantgdb.org/) and the National Center for Biotechnology Information (NCBI, www.ncbi.nlm.nih.gov). The BLAST E-value is believed to denote significant sequence similarity at a cut off value of 1×10^−8^.

QPCR results from flowers collected 24 h before and after anthesis suggest the existence of an opposite transcriptional pattern between Hawaiian and L1 lines (Figure 2). Relative expression of all loci is comparatively lower in L1, and unlike in Hawaiian papaya, the expression of all loci shows a sharp reduction after anthesis (Figure 2).

**Figure 2.**
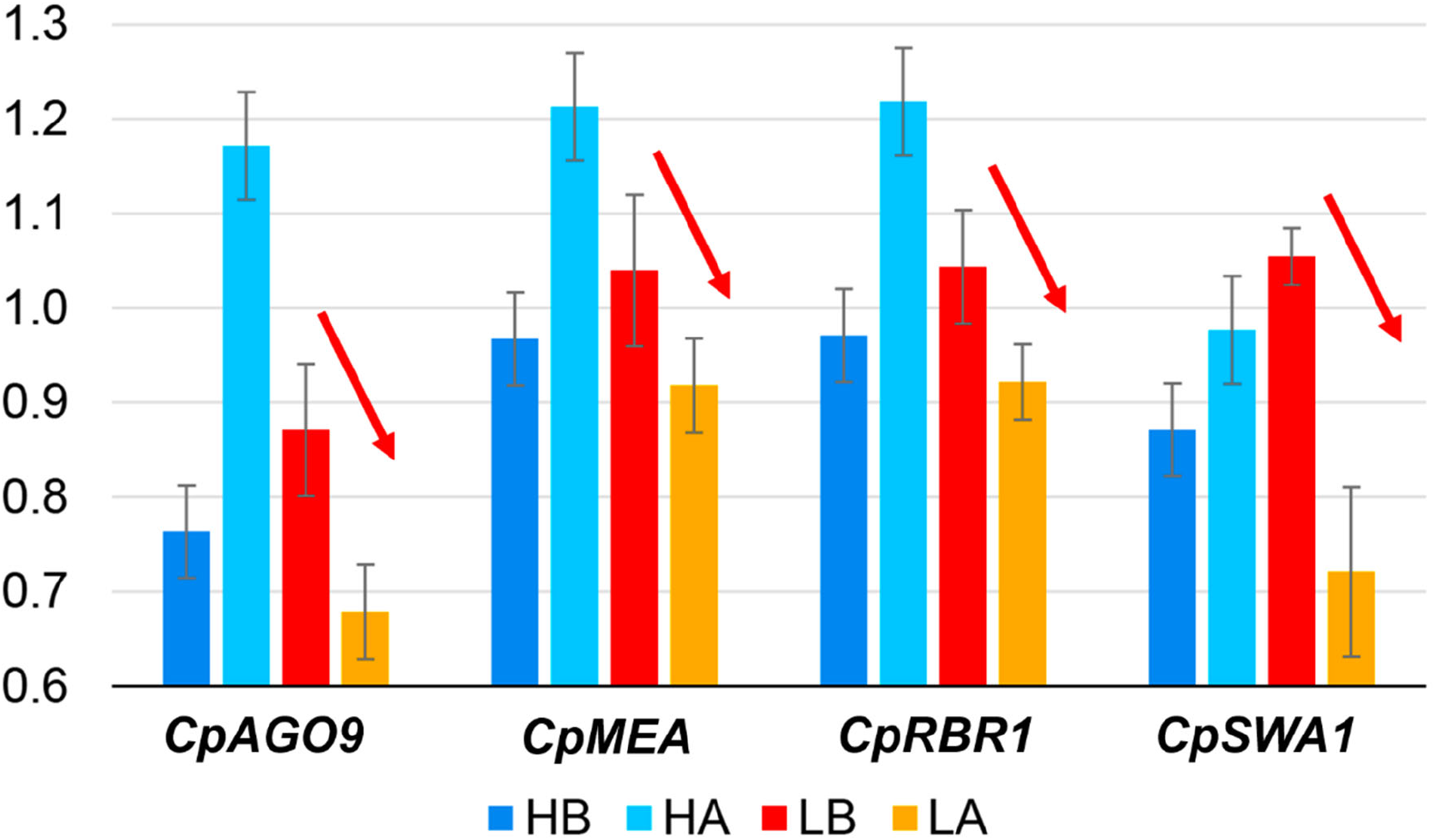
Transcriptional patterns in putative papaya regulators of embryo sac development. Results suggest differential expression between lines L1 (L) and Hawaiian (H). As compared with Hawaiian papaya, in the suspected apomictic line L1, the expression of loci *CpAGO9, CpMEA, CpRBR1* and *CpSWA1* appears to show a relative reduction in flowers both before anthesis (LB: L1 line before anthesis) and after anthesis (LA, L1 line after anthesis). The *Arabidopsis* homologs of these putative loci have been shown to repress endosperm (*AGO9, RBR1*) and embryo development (*MEA*), thus a reduction in expression might be interpreted as suggesting derepressed growth and autonomous egg and central cell differentiation. The *Arabidopsis* gene *SWA1* is believed to control female gametophyte fertility and central cell proliferation, and its putative papaya homolog (*CpSWA1*) showed reduced expression in L1, suggesting deregulated proliferation. The expression pattern of all loci in line Hawaiian is opposite to L1 both before anthesis (HB) and after anthesis (HA). qPCR experiments were performed in triplicate from three biological samples.

Work in *Arabidopsis* indicates that loss in the activity of the RNA-dependent DNA methylation (RdDM) gene *ARGONAUTE9* (*AGO9*) causes differentiation of ectopic cells in the embryo sac (Hernández-Lagana *et al*., 2016). Dominant mutations in *AGO9* lead to phenotypes in which somatic sporophytic cells give rise to a female gametophyte without undergoing meiosis (Rodríguez-Leal and Vielle-Calzada, 2012). In general, mutations in genes from the RdDM pathway may lead to similar phenotypes, which suggests that silencing of heterochromatic repetitive sequences is crucial to differentiate between sexual and apomictic cell fate (Rodríguez-Leal and Vielle-Calzada, 2012). *Arabidopsis* genes of the Polycomb Group complexes (PcG) mediate the transition from vegetative to reproductive development (Holec and Berger, 2012) and enforce transcriptional repression by catalyzing H3K27 trimethylation and H2A ubiquitination (Wang *et al*., 2016). For instance, mutants of *MEDEA* (*MEA*) show endosperm proliferation without fertilization (Koltunow and Grossniklaus, 2003) and suppression of central cell proliferation (Kiyosue *et al*., 1999). MEDEA is a SET-domain PcG protein that may affect the degree of localized chromatin condensation, gene transcription and cell proliferation (Kiyosue *et al*., 1999).

The *Arabidopsis RETINOBLASTOMA RELATED1* (*RBR1*) gene is the direct repressor of the stem cell factor WUSCHEL (WUS) and PROLIFERATING CELL NUCLEAR ANTIGEN 1 (PCNA1) (Zhao *et al*., 2017). Besides its main role in regulating cell cycle progression, RBR1 also cooperates with the PcG complex PRC2, which includes MEA, and has an important role in controlling gametophyte development (Kuwabara and Gruissem, 2014). In *rbr1* mutants, meiocytes undergo several mitotic divisions, resulting in extra meiocytes, which develop into megaspores that can successfully attract a pollen tube and be fertilized but do not develop (Zhao *et al*., 2017). Finally, the *slow walker1* mutation can cause partial female sterility because of delayed development of the female gametophyte at anthesis (Yang *et al*., 2010). *Arabidopsis SLOW WALKER-1* (*SWA1*) encodes a nucleolar WD40-containing protein involved in the processing of pre-18S rRNA (Yang *et al*., 2010). *SLOW WALKER*-mediated RNA processing and ribosome biogenesis may be important for the progression of mitotic division cycles during *Arabidopsis* embryo-sac development (Liu *et al*., 2010).

Taken together, reduced expression of the putative papaya homologs for *AGO9, MEA, RBR1* and *SWA1* in L1 embryo sacs (Figure 2) may account for sporophytic tissue proliferation in L1 embryo sacs. Although we do not know what causes these changes in the transcriptional patterns of L1, *AGO9* has a prominent role in binding heterochromatic small interfering RNAs (siRNAs) that mostly target repetitive genomic regions and transposable elements (Hernández-Lagana *et al*., 2016). Thus, these changes in transcriptional activity may occur due to generalized failure to perform RNA-directed DNA methylation. In *Arabidopsis*, the patterns of transcriptional regulation and protein localization of AGO9 in developing ovules vary between ecotypes (Rodríguez-Leal *et al*., 2015).

In Venezuela, Vegas *et al*. (2003) reported that one could harvest fruits from covered female flowers belonging to the papaya cultivars Costa Rica, Larga, Paraguanera, and Solo 6, 7 and 8 at a rate of 32% to 81% (Vega *et al*., 2003). Fruits carry seeds at a rate of 64%, and these seeds do not develop endosperm but show seemingly functional embryos (Vega *et al*., 2003). Embryos from the cultivar Larga were cultured *in vitro*, and somatic embryogenesis and subsequent plantlet regeneration were possible (Vega *et al*., 2003). These plantlets were grown in the field and were apparently identical in morphology (Vega *et al*., 2003). Multiple embryos were observed at a low rate of 5/3000 per dissected seeds (Vega *et al*., 2003).

## Conclusions

Dual staining of tissue sections is a conventional method that has proved successful in plant cytological studies (Blokhina *et al*., 2017). Our results indicate that this methodology may allow for identification of most features in the papaya ovules. In both papaya lines we could identify the contour of antipodal cells, polar nuclei/central cell, synergids and egg from about 10-15 samples. Deep coloring by the membrane stain Alcian blue suggested that their possible location might be similar to that observed in *A. thaliana* (Sundaresan and Alandete-Saez, 2010), with the egg cell placed right behind the synergids, and a large vacuolated central cell placed behind the egg cell (Johnson *et al*., 2019). In fact, large vacuolar membranes are associated with all these cells (Shi and Yang, 2011). Papaya ovules conform to a *Polygonum*-type architecture, but significant variation in morphology was observed. Although we assume that papaya ovule development is synchronic within a flower, synchronicity has not been determined experimentally yet.

Our results also hint at the existence of asexual embryo development in papaya, as previously reported by Vega *et al*. (2003). Variation in transcriptional patterns was also observed between different papaya parental lines at the ovule level. A possible link may exist between apomictic development in ovules of L1 and downregulation of papaya homologs for *AGO9* (RdDM pathway), *MEA* (PcG-mediated histone remodeling), *RBR1* (cell cycle and histone remodeling) and *SWA1* (rRNA processing and cell cycle). The upstream determinant for such differences is unknown, and in-depth analyses of transcriptional activity may be needed to corroborate this very tentative hypothesis. However, breeding of interspecific hybrids between *Carica* and the related genera *Cylicomorpha, Horovitzia, Jacaratia, Jarilla*, and *Vasconcellea* often leads to the formation of parthenocarpic fruit (Fuentes and Santamaría, 2014), which suggests that perhaps hybridization and polyploidization events in papaya might lead to a situation of genomic shock characterized by dysregulation of gene expression, high gene dosage sensitivity and disturbed cell divisions, as reported often for other plant apomictics (Hojsgaard, 2018).

The large size of ovules in papaya suggests that because of its favorable cytology, this organism may well become an important emerging model system. Functional genomics of the reproductive apparatus of papaya may contribute to the advancement of plant breeding efforts in other species. For instance, in *Arabidopsis* and *Torenia*, a better understanding of pollen tube guidance within the embryo sac has allowed for the identification of cysteine-rich peptides from the defensin family, called LURE1 and LURE2, that work as ovule attractants to facilitate fertilization and fruit formation (Higashiyama and Takeuchi, 2015). Development of fluorescent cell markers for the egg cell and central cell have also allowed for better understanding synergid and antipodal cell development, for identifying novel regulators of *Arabidopsis* ovule development (Liu *et al*., 2017). Taken together further in-depth transcriptomic, cytological, and perhaps flow cytometry analyses in papaya ovules may be valuable from a scientific and agricultural point of view.

## Acknowledgements

This project was kindly funded by Vicerrectoría de Investigación (University of Costa Rica) grant #B5A13. We thank Eric Mora-Newcomer (Fabio Baudrit Agricultural Research Station, University of Costa Rica) for kindly donating samples from his papaya breeding program, and to all students who contributed with their work. This manuscript was kindly edited by Ms. Laura Smales (BioMedEditing, Toronto, Canada). Pablo is a young member affiliate of TWAS/UNESCO and a member of the American Society of Plant Biologists (ASPB).

